# SATB homeobox 1 regulated genes in the mouse ectoplacental cone are important for placental development

**DOI:** 10.1101/2020.09.13.295584

**Authors:** V. Praveen Chakravarthi, Shaon Borosha, Anamika Ratri, Subhra Ghosh, Michael W. Wolfe, M. A. Karim Rumi

**Affiliations:** Department of Pathology and Laboratory Medicine, University of Kansas Medical Center, Kansas City, KS 66160; Department of Molecular and Integrative Physiology, University of Kansas Medical Center, Kansas City, KS 66160; Institute for Reproduction and Perinatal Research, University of Kansas Medical Center, Kansas City, KS 66160

**Author notes:** **Corresponding author:** M. A. Karim Rumi or V. Praveen Chakravathi, Institute for Reproduction and Perinatal Research, University of Kansas Medical Center, Kansas City, KS 66160. or.

**Keywords:** SATB homeobox 1, Transcriptome analysis, Mouse ectoplacental cone, Trophoblast cells, Placental development

## Abstract

SATB homeobox 1 (SATB1) is abundantly expressed in the stem-state of trophoblast cells but downregulated during trophoblast differentiation. It is also expressed in high levels in the mouse ectoplacental cones (EPCs). We detected that SATB1 is involved in maintaining the self-renewal of trophoblast stem cells and inhibiting trophoblast differentiation. In this study, we have identified SATB1-regulated genes in the mouse EPC and analyzed their potential functions. A total of 1618 differentially expressed genes were identified in *Satb1^null^* EPCs by mRNA sequencing. Remarkably 90% of the differentially expressed genes were found to be upregulated in *Satb1^null^* EPCs suggesting a transcriptional repressor role of SATB1 in mouse trophoblast cells. Ingenuity Pathway Analyses demonstrated that the differentially expressed genes in *Satb1^null^* EPCs are particularly linked to WNT and TGFβ signaling pathways, which regulate self-renewal of stem cells and cell differentiation. Moreover, twenty-six of the EPC genes that are known to be involved in placental development including *Eomes, Epas1, Fgfr2, Cdkn1c*, and *Plac9* were found dysregulated in *Satb1^null^* EPCs due to the loss of SATB1 expression. These genes are particularly involved in the formation of labyrinthine zone. Our results emphasize that SATB1-regulated genes in the mouse EPC contribute to key roles in the regulation of trophoblast differentiation and placental development.

## 1. INTRODUCTION

Special AT-rich sequence binding (SATB) proteins (SATB1 and SATB2) function as chromatin organizers and transcriptional regulators^[1–8]^. These homeobox proteins have been shown to play key roles in developmental processes, such as T cell differentiation^[3, 4, 8–10]^, erythroid development^[11]^, craniofacial patterning^[12]^, osteoblast differentiation^[12]^, cortical neuron organization^[6, 13–15]^, embryonic stem (ES) cell differentiation^[16]^, and the regulation of hematopoietic stem cell self-renewal^[17]^. Recently, we have demonstrated that SATB proteins contribute to the regulation of the trophoblast lineage^[18]^. SATB proteins are preferentially expressed in the trophoblast cell stem-state and are downregulated during differentiation^[18]^. We observed that SATB proteins maintain the TS cell stem-state and inhibit trophoblast differentiation^[18]^.

SATB proteins bind to AT-rich elements in matrix attachment regions of actively transcribing DNA and interact with chromatin remodeling proteins as well as transcription factors to induce gene activation or gene repression^[2–8, 12, 19]^. SATB proteins can act as docking sites for several chromatin remodeling enzymes including ACF and ISWI and can also recruit co-repressors like HDACs or co-activators like HATs directly to promoters and regulatory elements^[3, 20, 21]^. We have observed that SATB1 and SATB2 are abundantly expressed in the developing placenta and may possess overlapping roles in placental development^[18]^. We detected a high level of *Satb1* gene expression in embryonic day 7.5 (E7.5) ectoplacental cones (EPCs), which decreases as the gestation period progresses. As SATB1 acts as a transcriptional regulator of gene expression and is abundantly expressed in EPCs, we studied the whole transcriptome in *Satb1^null^* EPCs and compared it with that of wildtype EPCs.

## 2. METHOLODOLOGY

### 2.1. Satb1 knockout mouse

The *Satb1*-mutant mouse model was generated by targeted deletion of exon 4 in the *Satb1* gene. Exon 4 floxed *Satb1* mouse model was obtained from Jax Mice. The floxed mice were crossed with Meox2-Cre line to delete exon 4 of *Satb1* gene (**Fig. 1E, F**). Heterozygous mutant mice were backcrossed to CD1 background for 7 generation and the phenotypes of *Satb1^null^* mutants were found to be identical to previously published *Satb1* knockout model^[9]^. Heterozygous *Satb1*-mutant males and females were setup for mating, and the day of mating plug detection was considered as embryonic day 0.5 (E0.5). All the procedures were approved by the University of Kansas Medical Center (KUMC) Animal Care and Use Committee.

**Figure 1.**
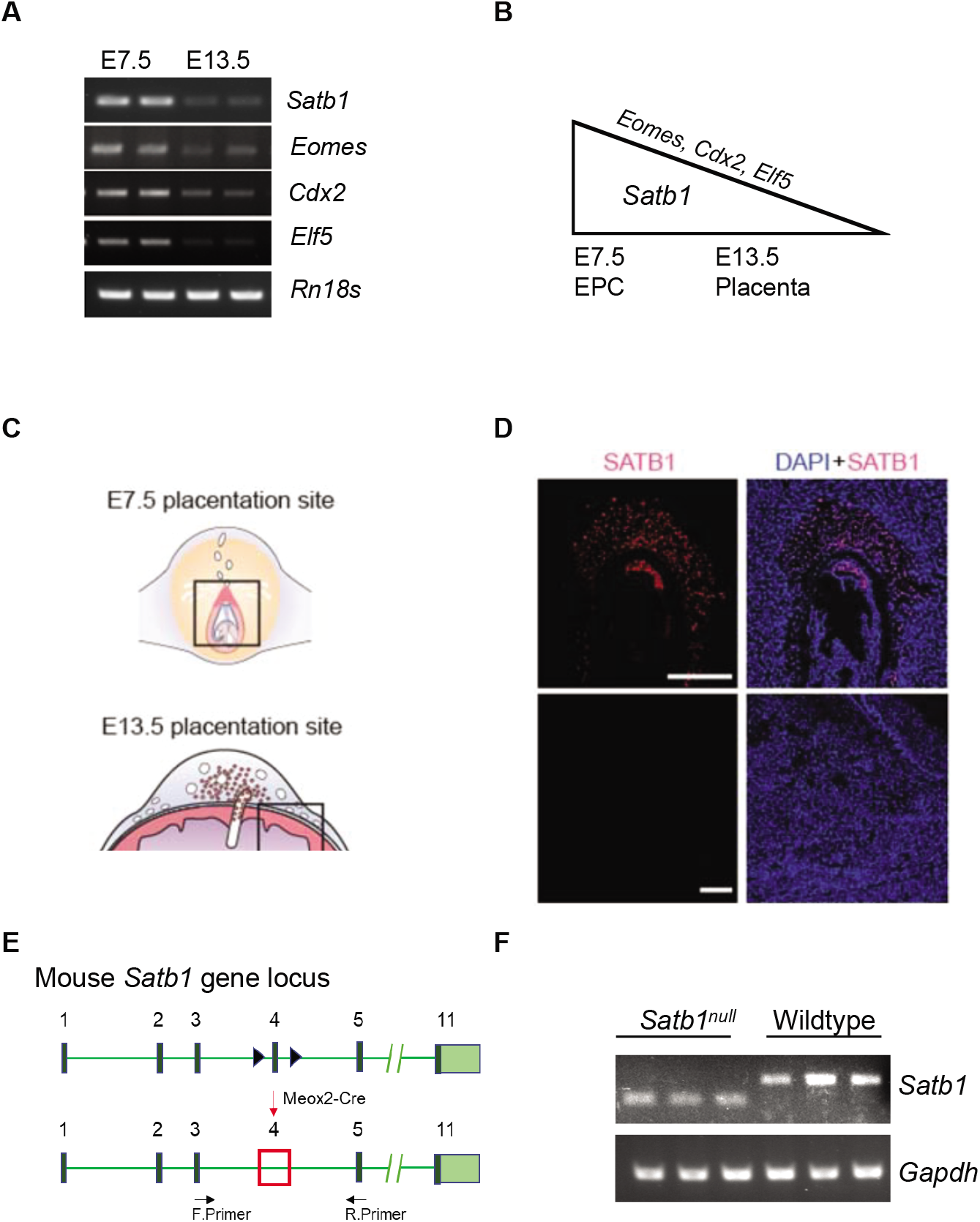
*Satb1* expression in mouse EPCs and generation of *Satb1^null^* mouse model. *Satb1* expression was determined at different stages of embryonic development i.e. on E7.5, and E13.5. Maximum expression of *Satb1* was found in E7.5 EPC as determined by RT-PCR (**A**). The expression of Satb1 correlated with that of *Eomes, Cdx2* and *Elf5* (A). Immunofluorescent staining of mouse E7.5 EPC and E13.5 placenta also demonstrated SATB1 expression in E7.5 EPC but not in E13.5 placenta (**C, D**). *Satb1*^null^ mouse was generated by targeted deletion of floxed exon 4 of mouse Satb1 gene by crossing with transgenic mouse expressing Meox2 Cre (**E, F**)).

### 2.2. Detection of SATB1 in mouse ectoplacental cone

Expression of *Satb1* was detected in mouse ectoplacental cones at mRNA and protein levels. For detection at mRNA level, pregnant *Satb1*-hetrozygous mutant females were euthanized on E7.5 to collect the conceptuses. Conseptuses were dissected under a stereomicroscope to isolate the ectoplacental cones, yolk sacs, and embryos. Genomic DNA extracted from the yolk sacs and embryos was used for geneotyping using the RED extract-N-Amp Tissue PCR Kit (Sigma-Aldrich, St. Louis, MO). Total RNA was purified from the ectoplacental cones using TRI-Reagent (Sigma-Aldrich) and cDNAs were prepared from 250 ng of total RNA pooled from multiple samples of same genotype was used to prepare cDNA using random primers and Superscript II reverse transcriptase (ThermoFisher Scientific). 0.5 μl of the 10x diluted cDNA and gene specific primer mix was used in a 10 μl qPCR reaction containing Power SYBR Green Master Mix (Applied Biosystems). RT-qPCR results were normalized to *Rn18s* expression and calculated by the comparative ΔΔCT method^[22, 23]^.

### 2.3. Immunofluorescent Microscopy

For SATB1 detection in EPC at protein levels, whole conceptuses were fixed in cold heptane and preserved at −80C freezer. Conceptuses were embedded in O.C.T. Compound (Fisher HealthCare, Houston, TX) and cryosection at 10μM thickness. Sections were preserved at −80C freezer until immunofluorescence staining. After blocking, the sections were incubated with appropriately diluted 129 primary antibodies: anti-SATB1 (ab109122, Abcam at 1:1000) at room temperature for 2h. After washing the 132 unbound primary antibodies, secondary antibody staining was performed with Alexa Fluor 568-133 or 488-labeled detection reagents (goat antirabbit, goat anti-mouse antibodies; Molecular Probes) at 1:200 dilution, and DNA staining was performed by DAPI (Prolong Gold Antifade Mounting Media, 135 Thermo Fisher Scientific). The images were captured on a Nikon Eclipse 80i microscope.

### 2.4. Sample collection, library preparation and RNA-sequencing

RNA quality was assessed by a Bioanalyzer at the KUMC Genomics Core, and samples with RIN values over 9 were selected for RNA-sequencing library preparation. RNA-seq libraries were prepared using the True-Seq mRNA kit (Illumina, San Diego, CA) as described previously^[24–26]^. RNA samples extracted from multiple wildtype or *Satb1^null^* ectoplacental cones were pooled to prepare each RNA-seq library. Approximately 500 ng of total RNA was used to prepare a RNA-seq library using the True-Seq mRNA kit (Illumina, San Diego, CA) as described previously^[24–26]^. The quality of RNA-seq libraries was evaluated by Agilent Analysis at the KUMC Genomics Core and the sequencing was performed on an Illumina HiSeq X sequencer (Novogene Corporation, Sacramento, CA).

### 2.5. RNA-seq data analyses

RNA-sequencing data were demultiplexed, trimmed, aligned, and analyzed using CLC Genomics Workbench 12.2 (Qiagen Bioinformatics, Germantown, MD) as described previously^[24–26]^. Through trimming, low-quality reads were removed, and good-quality reads were aligned with the *Mus musculus* genome (mm10) using default guidelines: (a) maximum number of allowable mismatches = 2, (b) minimum length and similarity fraction = 0.8, and (c) minimum number of hits per read = 10. Gene expression values were measured in transcripts per million (TPM). Differentially expressed genes were identified that had an absolute fold change of TPM ≥ 2 and a false discovery rate (FDR) *p*-value of ≤0.05.

### 2.6. Gene Ontology (GO) and disease pathway analyses for the RNA-sequencing data

Differentially expressed genes were subjected to Gene Ontology (GO) analysis (http://www.pantherdb.org) and categorized in biological, cellular and molecular function. Differentially expressed genes in *Satb1^null^* mouse TS cells were further analyzed by Ingenuity Pathway Analysis (IPA; Qiagen Bioinformatics, Germantown, MD) to build gene networks related to placental development. Functional analyses were performed towards understanding the biological pathways and functions altered in the *Satb1^null^* ectoplacental cone.

### 2.7. Validation of RNA-sequencing data

Differentially expressed genes were validated by RT-qPCR. RT-qPCR validation included cDNA samples prepared with wildtype and *Satb1^null^* EPC RNA. The genes were selected from the MGI data and IPA analyses that positively impacted the placental development. RT-qPCR primers for the detection of mouse EPC genes are shown in **Supplementary Table 1**.

### 2.7. Statistical analysis

Each RNA-seq library or cDNA was prepared from pooled RNA samples extracted from at least 6 different ectoplacental cones of the same genotype. Each group for RNA sequencing consisted of three independent libraries and the differentially expressed genes were identified by CLC Genomics workbench as described previously^[24–26]^. RT-qPCR validation included at least six cDNA samples prepared from wildtype or *Satb1^null^* EPC RNA. The experimental results are expressed as mean + standard error (SE). The RT-qPCR results were analyzed for one-way ANOVA, and the significance of mean differences was determined by Duncan’s *post doc* test, with *p* < 0.05. All the statistical calculations were done using SPSS 22 (IBM, Armonk, NY).

## 3. RESULTS

### 3.1. Expression of Satb1 in the mouse EPC

*Satb1* mRNA and protein were abundantly expressed in the embryonic day (E7.5) wildtype EPCs, but the expression decreased with the progression of gestation (**Fig. 1A-D**). The expression of proliferating trophoblast-specific transcription factors including *Eomes, Cdx2*, and *Elf5* also correlates with the expression of *Satb1* in the E.7.5 EPC and E13.5 placenta (**Fig. 1A, B**). The expression of SATB1 protein was confirmed by immunofluorescence staining, which detected it in the E7.5 EPC but not the E13.5 placenta (**Fig. 1C, D**). We generated a *Satb1^null^* mutant model by targeted deletion of exon 4 of the mouse *Satb1* gene (**Fig. 1E, F**).

### 3.2. RNA-sequencing data of wildtype and Satb1^null^ EPCs

Transcriptome data-sets were generated by sequencing mRNA purified from EPCs dissected out from E7.5 wildtype and *Satb1^null^* mouse conceptuses. The raw data have been deposited to NCBI SRA under PRJNA562732. SRR10032643, SRR10032642, and SRR10032641 represent transcriptome data obtained from the wildtype EPCs, while SRR10032640, SRR10032639, and SRR10032638 represent those from *Satb1^null^* EPCs. Analyzed data include the differentially expressed genes in *Satb1^null^* EPCs (**Supplementary Table 2 and 3**).

### 3.3. Differentially expressed genes in Satb1^null^ mouse EPCs

RNA-sequencing data were analyzed by using CLC Genomics Workbench to identify the differentially expressed genes in *Satb1^null^* EPCs. Of the total 25,749 reference genes in the mm10 genome, 17,005 genes were detected in wildtype and 17,570 in *Satb1^null^* EPCs. Analyses of the detected genes for level of gene expression revealed that ~28% had a very low abundance (<1 TPM), ~26% had low abundance (1-10 TPM), ~37% had high abundance (10-100 TPM), and only ~9% of the genes had a very high abundance (> 100 TPM) (**Fig. 2A**). Among these genes, 1618 were differentially expressed (absolute fold change ≥2, FDR *p*-value ≤ 0.05) in *Satb1^null^* EPCs, with 157 downregulated and 1461 upregulated (**Supplementary Table 2 and 3**). These differentially expressed genes are also evident in the hierarchical clustering and volcano plots, which demonstrate that most of the differentially expressed genes were upregulated in *Satb1^null^* EPCs (**Fig. 2B, C**). Based on fold changes in gene expression in *Satb1^null^* EPCs and maximum TPM values, top 50 downregulated and top 50 upregulated genes are listed in **Table 1 and 2**.

**Figure 2.**
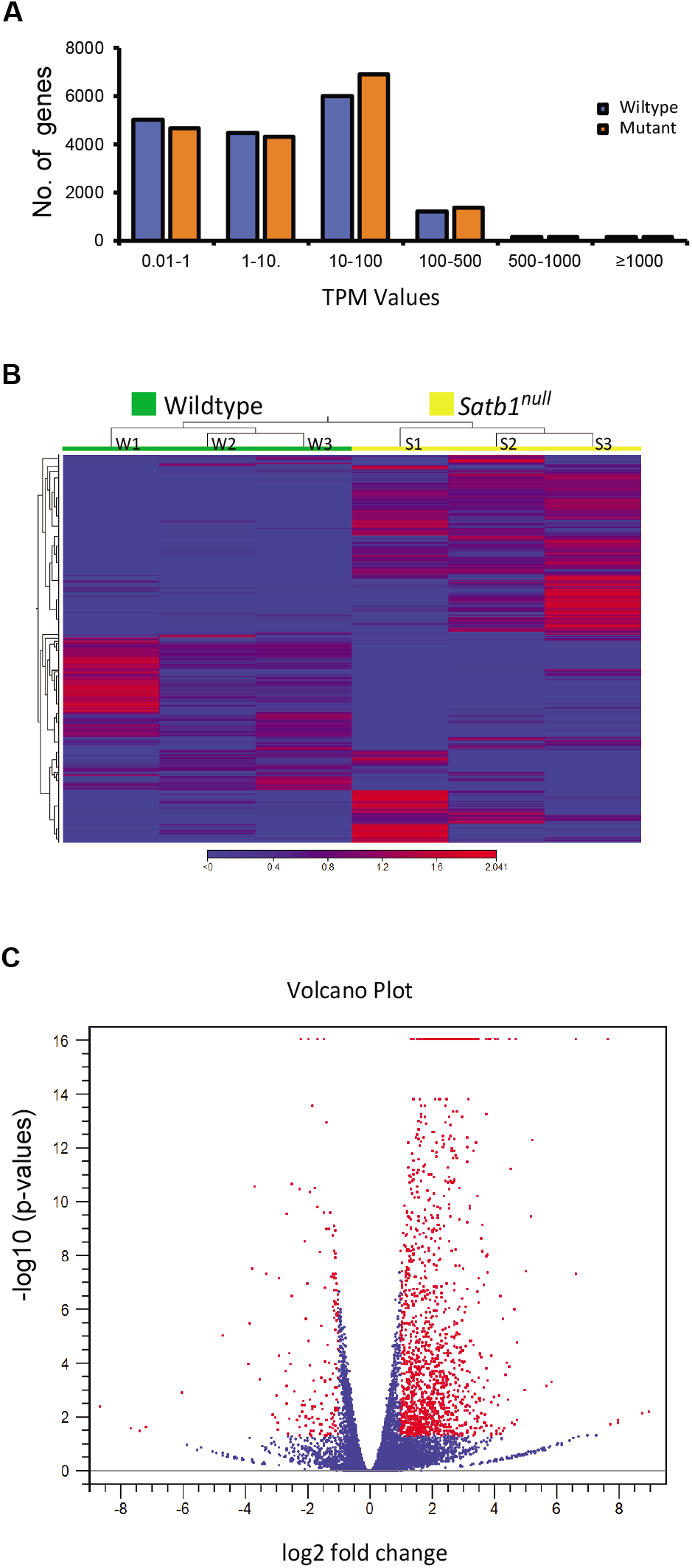
RNA sequencing analysis of wildtype and *Satb1^null^* EPC. EPCs along with yolksacs and embryos were isolated from E7.5 conceptuses collected preganant *Satb1* heterozygous mutant female mice crossed with *Satb1* heterozygous males. EPCs were used for RNA purification. Yolk sacs and embryos were used for genotyping. RNA was isolated from individual EPCs and pooled RNAs from mutltiple samples were used for library preparation and RNA sequencing. RNA-seq data were analyzed CLC Genomics Workbench and differentially expressed genes were classified into several groups based on their TPM values (**A**). Hierarchical clustering was also performed on the differentially expressed genes between wildtype and *Satb1^null^* EPCs (**B**). Volcano plot between wildtype and *Satb1^null^* EPC showed a range of differentially expressed genes at different fold changes (**C**).

**Table 1.** Top 50 downregulated genes in *Satb1^null^* mouse ectoplacental cones.

| Name | Chromosome | Region | Max group mean | Log<sub>2</sub> fold change | Fold change | P-value | FDR p-value | Bonferroni | ENSEMBL |
| --- | --- | --- | --- | --- | --- | --- | --- | --- | --- |
| *mt-Co3* | MT | 8607..9390 | 9,259.87 | -1.24 | -2.36 | 3.19E-12 | 2.73E-10 | 8.20E-08 | ENSMUSG00000064358 |
| *mt-Nd1* | MT | 2751..3707 | 5,340.26 | -1.46 | -2.76 | 0 | 0 | 0 | ENSMUSG00000064341 |
| *mt-Nd3* | MT | 9459..9806 | 1,768.53 | -1.67 | -3.18 | 0 | 0 | 0 | ENSMUSG00000064360 |
| *Hbb-bs* | 7 | complement(103826534..103828096) | 1,517.21 | -1.81 | -3.52 | 2.22E-16 | 3.04E-14 | 5.72E-12 | ENSMUSG00000052305 |
| *Hba-a1* | 11 | 32283511..32284465 | 1,324.87 | -1.48 | -2.78 | 3.14E-12 | 2.70E-10 | 8.10E-08 | ENSMUSG00000069919 |
| *Gm10260* | 13 | complement(97760072..97760632) | 1,072.62 | -2.51 | -5.71 | 4.20E-06 | 1.20E-04 | 0.11 | ENSMUSG00000069117 |
| *Hmox1* | 8 | 75093621..75100589 | 753.27 | -1.37 | -2.58 | 1.50E-11 | 1.16E-09 | 3.85E-07 | ENSMUSG00000005413 |
| *Prl3d2* | 13 | 27121698..27127555 | 578.43 | -1.74 | -3.35 | 3.58E-13 | 3.39E-11 | 9.22E-09 | ENSMUSG00000062737 |
| *Gm11361* | 13 | 28257528..28258134 | 463.46 | -2.56 | -5.9 | 1.41E-06 | 4.62E-05 | 0.04 | ENSMUSG000000113061 |
| *Lars2* | 9 | 123366927..123462666 | 421.26 | -1.3 | -2.47 | 3.52E-06 | 1.03E-04 | 0.09 | ENSMUSG00000035202 |
| *Hspe1* | 1 | 55088132..55091307 | 413.75 | -1.2 | -2.29 | 1.13E-09 | 6.75E-08 | 2.91E-05 | ENSMUSG00000073676 |
| *Serpinb9e* | 13 | 33249612..33260850 | 323.92 | -1.31 | -2.47 | 1.45E-11 | 1.13E-09 | 3.74E-07 | ENSMUSG00000062342 |
| *Prl3d3* | 13 | 27156790..27162593 | 272.24 | -1.97 | -3.92 | 0 | 0 | 0 | ENSMUSG00000062201 |
| *Gstp1* | 19 | complement(4035407..4037985) | 261.69 | -1.39 | -2.62 | 8.88E-16 | 1.14E-13 | 2.29E-11 | ENSMUSG00000060803 |
| *Prl6a1* | 13 | 27312627..27319252 | 199.42 | -1.26 | -2.39 | 1.72E-05 | 4.07E-04 | 0.44 | ENSMUSG00000069259 |
| *Hbb-bt* | 7 | complement(103812524..103813996) | 177.54 | -1.91 | -3.75 | 4.72E-13 | 4.39E-11 | 1.22E-08 | ENSMUSG00000073940 |
| *mt-Nd6* | MT | complement(13552..14070) | 177.54 | -1.58 | -2.99 | 1.22E-10 | 8.38E-09 | 3.15E-06 | ENSMUSG00000064368 |
| *Tpm3* | 3 | 90072649..90100902 | 146.79 | -1.43 | -2.7 | 3.10E-09 | 1.74E-07 | 7.97E-05 | ENSMUSG00000027940 |
| *Ahcy* | 2 | complement(155059310..155074497) | 86.42 | -2.89 | -7.41 | 1.23E-09 | 7.27E-08 | 3.16E-05 | ENSMUSG00000027597 |
| *Utf1* | 7 | 139943789..139945112 | 65.71 | -1.19 | -2.29 | 2.72E-11 | 2.04E-09 | 7.01E-07 | ENSMUSG00000047751 |
| *S100g* | X | complement(162961992..162964599) | 49.38 | -1.77 | -3.41 | 6.35E-05 | 1.24E-03 | 1 | ENSMUSG00000040808 |
| *Prl7c1* | 13 | complement(27773495..27780804) | 37.72 | -2.21 | -4.61 | 0 | 0 | 0 | ENSMUSG00000060738 |
| *Gm7879* | 5 | 43913144..43913917 | 26.34 | -1.35 | -2.55 | 6.52E-07 | 2.33E-05 | 0.02 | ENSMUSG00000033036 |
| *Tpbpa* | 13 | complement(60938492..60941984) | 23.01 | -1.66 | -3.16 | 1.94E-12 | 1.71E-10 | 5.00E-08 | ENSMUSG00000033834 |
| *Pfdn4* | 2 | 170496428..170519123 | 21.25 | -1.18 | -2.26 | 1.82E-08 | 8.87E-07 | 4.69E-04 | ENSMUSG00000052033 |
| *Mcrip2* | 17 | complement(25863698..25868764) | 16.49 | -1.19 | -2.28 | 7.86E-05 | 1.49E-03 | 1 | ENSMUSG00000025732 |
| *Fam213b* | 4 | complement(154895504..154899135) | 13.72 | -1.2 | -2.3 | 1.16E-03 | 0.01 | 1 | ENSMUSG00000029059 |
| *Zfp979* | 4 | complement(147611937..147642513) | 13.55 | -2.7 | -6.48 | 7.86E-06 | 2.07E-04 | 0.2 | ENSMUSG00000066000 |
| *Prl2b1* | 13 | complement(27383339..27390846) | 13.08 | -1.9 | -3.72 | 4.33E-06 | 1.24E-04 | 0.11 | ENSMUSG00000069258 |
| *Gm20075* | 13 | complement(96079216..96081376) | 12.09 | -1.97 | -3.93 | 4.18E-07 | 1.56E-05 | 0.01 | ENSMUSG000000114133 |
| *Cmtm5* | 14 | 54936261..54939274 | 11.16 | -1.38 | -2.6 | 9.57E-05 | 1.76E-03 | 1 | ENSMUSG00000040759 |
| *Alas2* | X | 150547375..150570638 | 10.76 | -1.53 | -2.88 | 1.02E-07 | 4.34E-06 | 2.63E-03 | ENSMUSG00000025270 |
| *Gm9513* | 9 | 36475638..36477186 | 10.73 | -1.92 | -3.78 | 1.18E-03 | 0.01 | 1 | ENSMUSG00000090710 |
| *Ccnb1ip1* | 14 | complement(50789248..50807732) | 10.54 | -2.08 | -4.23 | 4.28E-11 | 3.13E-09 | 1.10E-06 | ENSMUSG00000071470 |
| *Arg2* | 12 | 79130777..79156301 | 10.47 | -1.24 | -2.37 | 2.65E-03 | 0.02 | 1 | ENSMUSG00000021125 |
| *Tfpi2* | 6 | complement(3962595..3988919) | 9.11 | -1.98 | -3.95 | 1.97E-09 | 1.14E-07 | 5.06E-05 | ENSMUSG00000029664 |
| *Gm10031* | 1 | 156524012..156526664 | 8.45 | -1.52 | -2.88 | 1.58E-05 | 3.79E-04 | 0.41 | ENSMUSG000000101523 |
| *Car12* | 9 | 66713686..66766845 | 7.96 | -1.41 | -2.66 | 3.21E-06 | 9.55E-05 | 0.08 | ENSMUSG00000032373 |
| *Ndufaf2* | 13 | complement(108002715..108158623) | 7.47 | -1.18 | -2.26 | 1.02E-05 | 2.60E-04 | 0.26 | ENSMUSG00000068184 |
| *Dynlt1c* | 17 | 6601671..6609774 | 7.05 | -2.24 | -4.74 | 3.97E-13 | 3.71E-11 | 1.02E-08 | ENSMUSG00000000579 |
| *Zfp985* | 4 | 147553277..147585198 | 7.02 | -2.66 | -6.3 | 3.66E-12 | 3.11E-10 | 9.42E-08 | ENSMUSG00000065999 |
| *Tmem59l* | 8 | complement(70483867..70487358) | 6.91 | -2.5 | -5.65 | 6.81E-09 | 3.54E-07 | 1.75E-04 | ENSMUSG00000035964 |
| *Gm13698* | 2 | 83955693..83958185 | 6.6 | -1.6 | -3.04 | 6.12E-03 | 0.05 | 1 | ENSMUSG00000094125 |
| *Krt14* | 11 | complement(100203162..100207548) | 5.94 | -1.81 | -3.51 | 1.92E-05 | 4.48E-04 | 0.5 | ENSMUSG00000045545 |
| *Rpl39l* | 16 | 10170226..10174911 | 5.81 | -1.38 | -2.6 | 6.24E-03 | 0.05 | 1 | ENSMUSG00000039209 |
| *Sult4a1* | 15 | complement(84076097..84105754) | 5.65 | -1.41 | -2.65 | 2.33E-05 | 5.28E-04 | 0.6 | ENSMUSG00000018865 |
| *Rac3* | 11 | 120721470..120723969 | 5.33 | -1.24 | -2.36 | 5.89E-03 | 0.05 | 1 | ENSMUSG00000018012 |
| *Tmem125* | 4 | complement(118540941..118544044) | 5.19 | -1.2 | -2.3 | 5.71E-04 | 7.50E-03 | 1 | ENSMUSG00000050854 |
| *Il17d* | 14 | 57524777..57543166 | 5.08 | -3.29 | -9.81 | 8.74E-10 | 5.32E-08 | 2.25E-05 | ENSMUSG00000050222 |
| *Fam83a* | 15 | 57985419..58011009 | 5.07 | -3.66 | -12.65 | 3.00E-13 | 2.87E-11 | 7.73E-09 | ENSMUSG00000051225 |

**Table 2.**
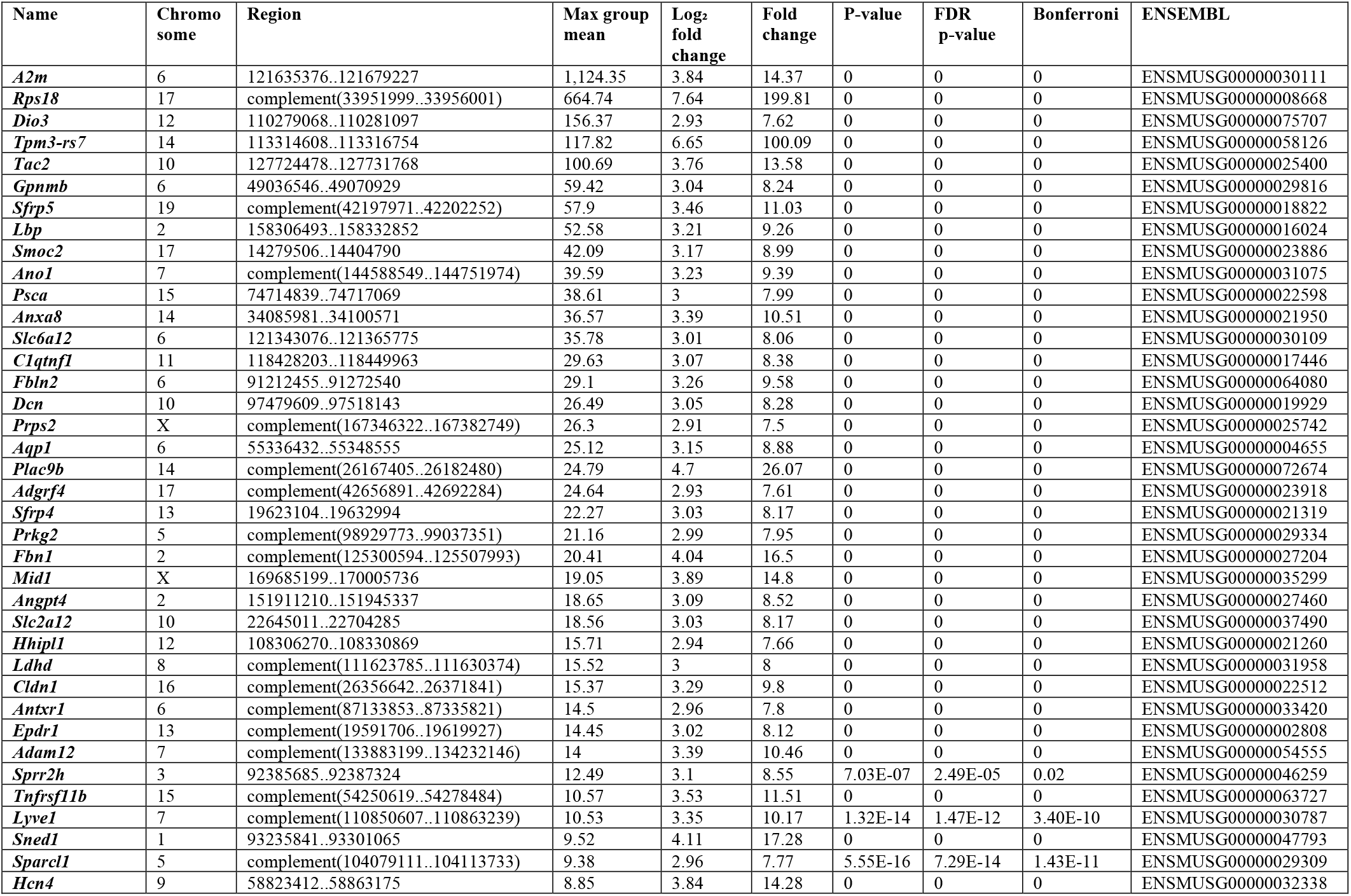
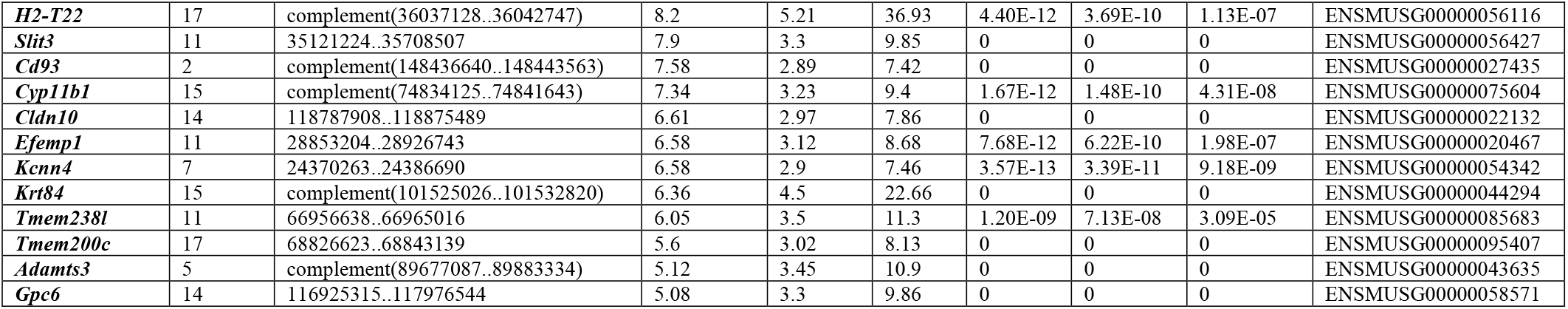
Top 50 upregulated genes in *Satb1^null^* mouse ectoplacental cones.

### 3.4. Gene Ontology (GO) analyses of the differentially expressed genes

GO analysis classified the differentially expressed genes into three categories: Biological process (**Fig. 3A**), Molecular function (**Fig. 3B**) and Cellular component (**Fig. 3C**). GO analysis revealed that the majority of the genes in the biological process group were involved in cellular processes, biological regulation, or cell signaling (**Fig. 3A**). The genes in molecular function were involved in protein-protein interactions, catalytic activity, and molecular and transcriptional regulation (**Fig. 3B**). The genes in cellular component were predominantly involved in cell parts, membranes, organelles, and protein-protein complexes (**Fig. 3C**).

**Figure. 3.**
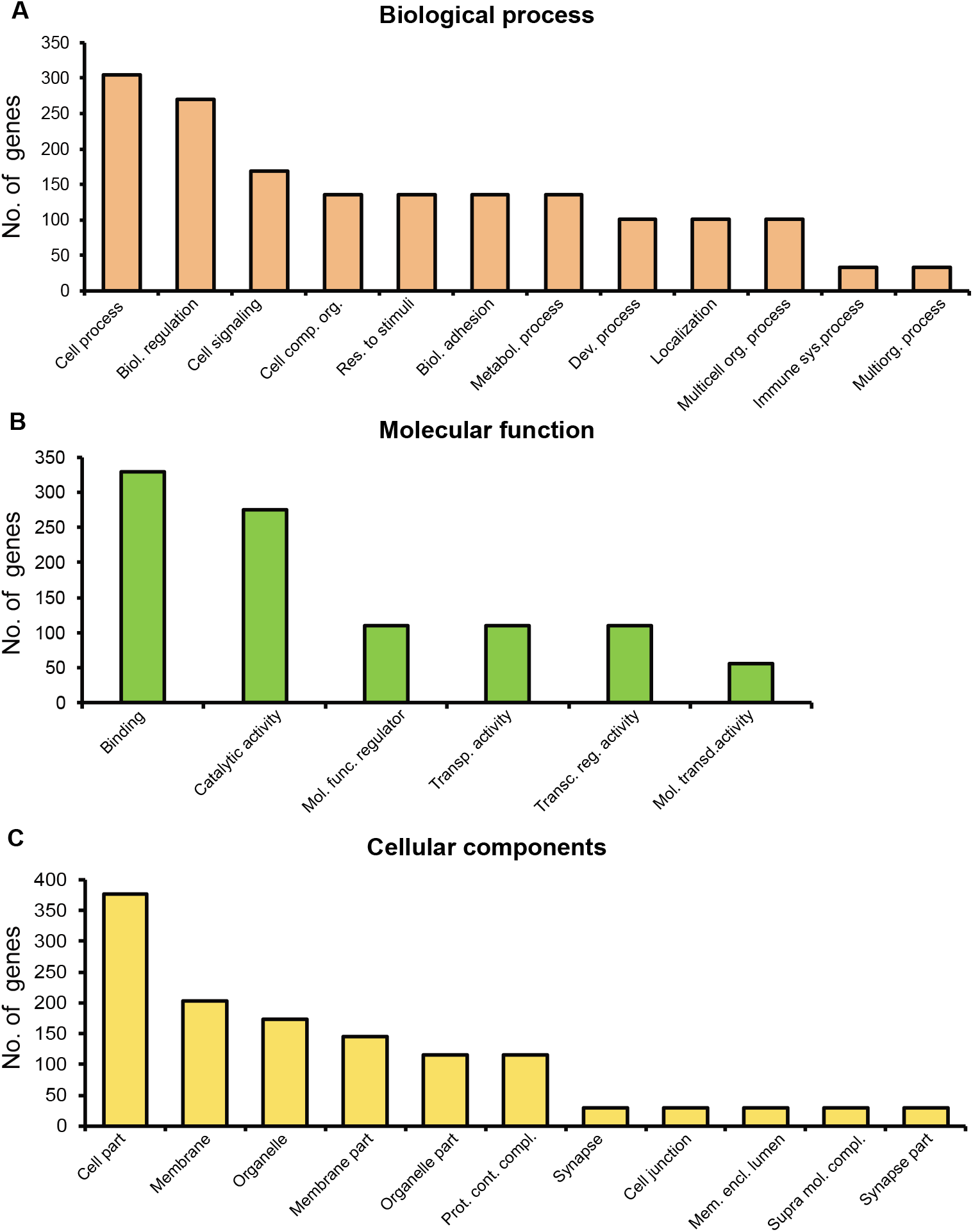
Classification of differentially expressed genes based on Gene Ontology. The differentially expressed genes in *Satb1^null^* EPCs were subjected to Panther Classification Analysis (http://pantherdb.org). Differentially expressed genes were classified based on Biological process (A), Molecular function (B) and Cellular components (C).

### 3.5. Ingenuity Pathway Analysis (IPA) of the differentially expressed genes

IPA of the differentially expressed genes in *Satb1^null^* EPCs identified upregulation of genes related to decreased pluripotency, increased differentiation, and development of the trophectoderm, endoderm and mesoderm (**Supplementary Fig.1**). The pathway analyses also detected downregulated genes in *Satb1^null^* EPCs are required for pluripotency and self-renewal of stem cells (**Supplementary Fig.1**). The expression of selected genes involved in the WNT Pathway including *Wnt2, Wnt3a, wnt10a*, and *Wnt10b* that showed significant upregulation was verified by RT-qPCR (**Fig. 4A-D**). Similarly, marked upregulation of several genes involved in the BMP and TGFβ pathways including *Bmp2, Bmp8, Smad1, Smad9*, and *Tgfbr2* (**Fig. 4E-I**) was also confirmed by RT-qPCR analyses. Finally, we verified the downregulation of *Utf1* in the *Satb1^null^* EPC, which is crucial for the self-renewal of stem cells (**Fig. 4J**).

**Figure 4.**
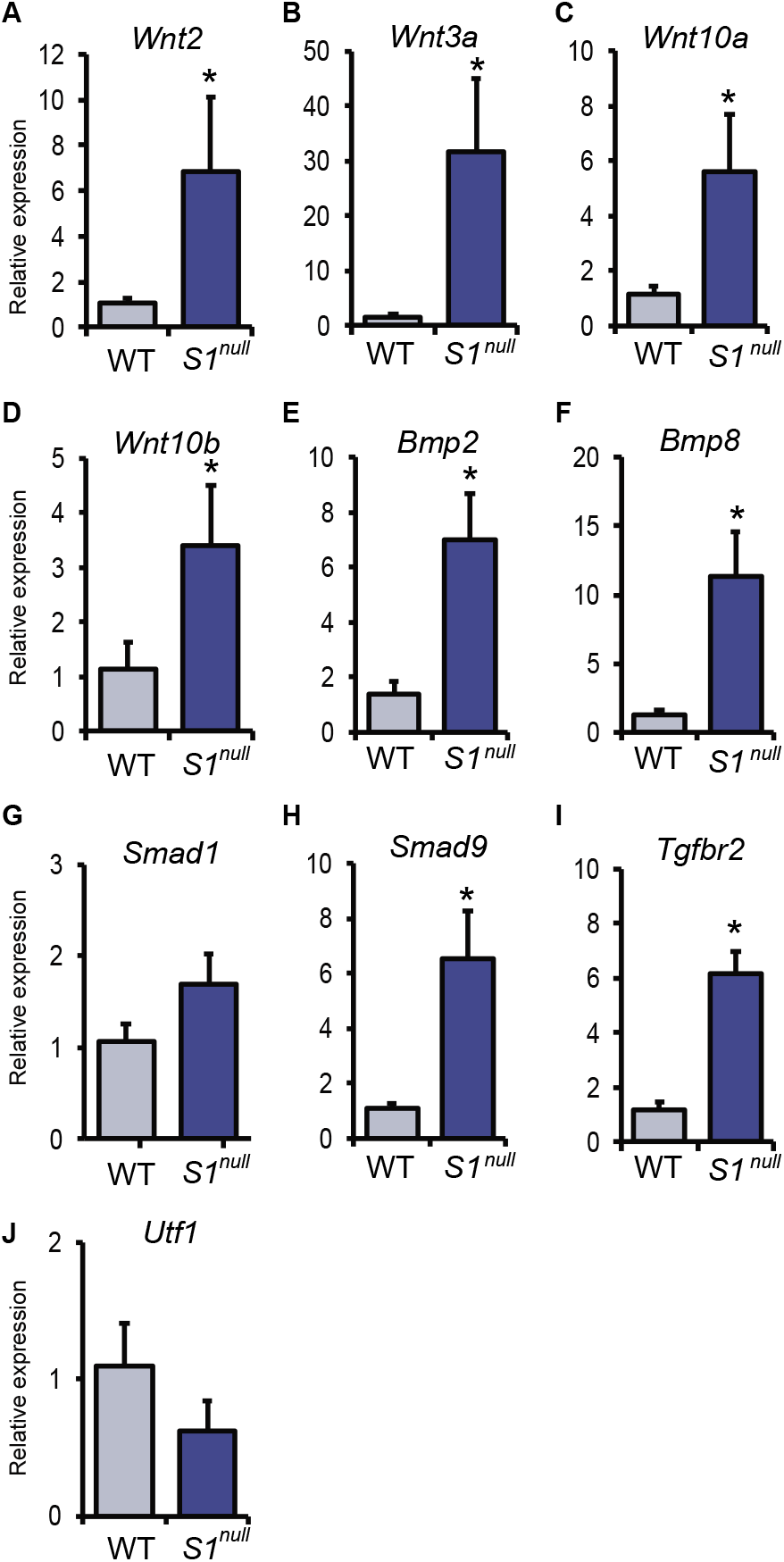
Differentially expressed genes involved in stem cell pluripotency. Differentially expressed genes in *Satb1^null^* EPCs were subjected to IPA analysis and identified a subset of genes involved in stem cell pluripotency **(Supplementary Figure 1)**. We further identified the upregulation of genes involved in WNT, and TGFβ pathway that can lead to abnormal differentiation of trophoblast and defective placental development. The expression of selected genes in the WNT (**A-D**), and TGFβ pathway (**E-I**) were further confirmed by RT-qPCR analyses.

### 3.6. Differentially expressed genes involved in placental development

Differentially expressed genes in *Satb1^null^* EPCs were compared with a list of genes curated from the MGI database (www.informatics.jax.or) that have been known to be involved in placental development. This comparison identified a large number of genes common to both groups, which were subjected to IPA. IPA analysis generated a gene network that connected differentially expressed genes to the formation of the placenta, and the morphology of the placenta, particularly the labyrinthine zone (**Fig. 5A**). Several genes were selected from this group on the basis of relative fold changes and TPM values in RNA-seq data and their differential expressions were further confirmed by RT-qPCR (**Fig. 5B-F**). While *Adamts3, Mmp14, Calcrl*, and *Syna* were significantly upregulated (**Fig. 5B-E**), expression of *Utf1* and *Cdkn1c* were downregulated in the *Satb1^null^* EPCs (**Fig. 4K, Fig. 5F**).

**Figure 5.**
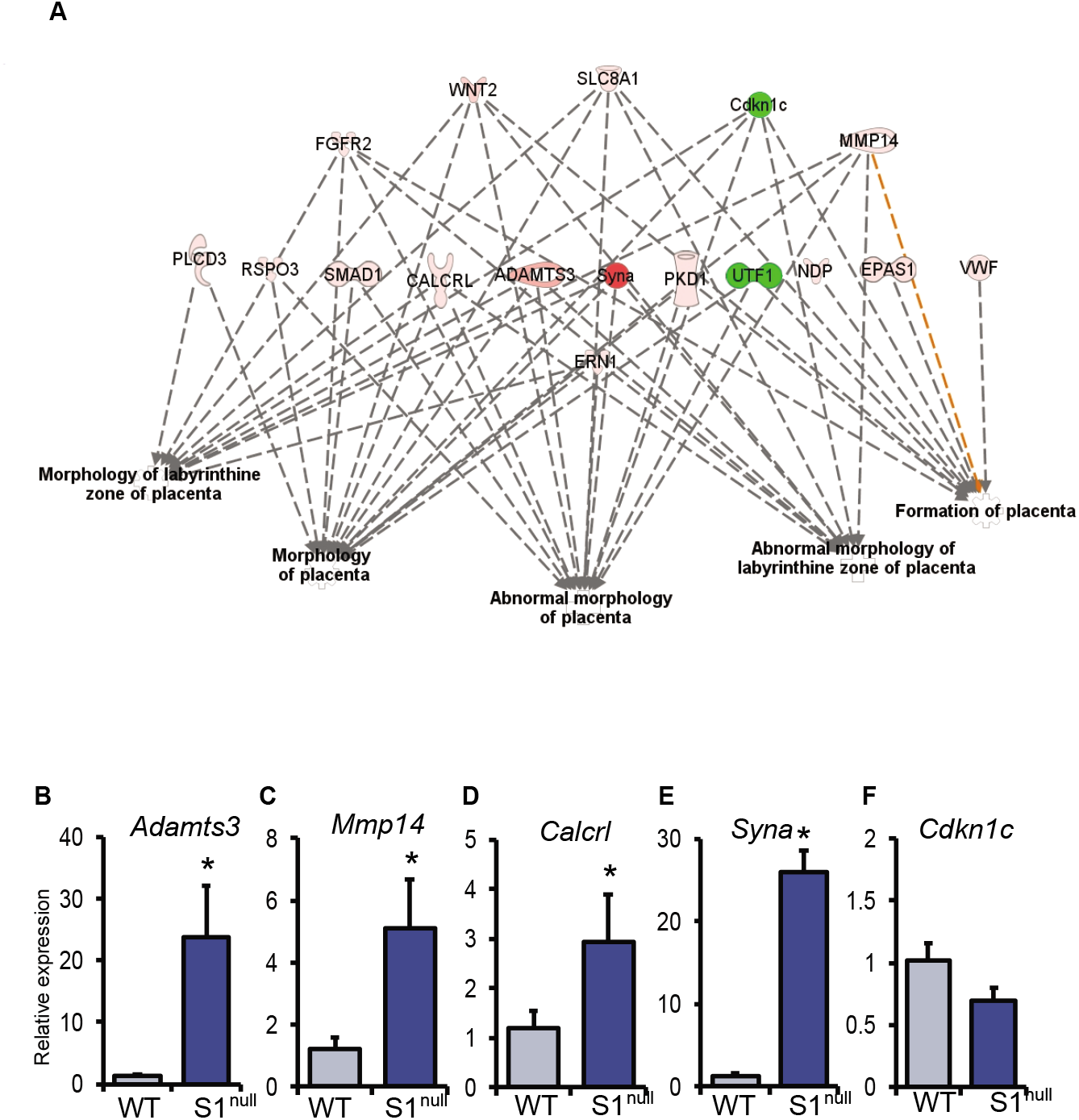
DEGs involved in placental development. Differentially expressed genes involved in placental development were screened using a subset of genes curated from MGI database (www.informatics.jax.or). These genes were subjected to IPA analysis and found a network of genes affected which leads to abnormal development of placenta (**A**). The expression of selected genes that can affect development of labyrinthine zone were further confirmed by RT-qPCR analyses (**B-F**).

## 4. DISCUSSION

SATB1 is expressed in high levels in mouse EPCs, suggesting a potential role for this chromatin organizer and transcriptional regulator in early placental development. This study analyzed the differentially expressed genes in *Satb1^null^* mouse EPCs during early embryonic development. RNA-seq datasets were used to identify SATB1-regulated genes in mouse EPCs and understand their potential role in placental development.

Previous studies have demonstrated that SATB proteins can either repress or activate gene expression in a tissue-specific manner^[3, 20, 21]^. Our RNA-seq data demonstrated that ~90% of the differentially expressed genes (1461 out of 1628) were upregulated in *Satb1^null^* EPCs due to the loss of SATB1 (Fig. 2), which clearly indicates that SATB1 acts as a suppressor of gene expression in EPCs. Notably, as per the TPM values, a significant proportion of these genes showed a high level of expression in mouse EPCs.

GO enrichment revealed that a significant number of the differentially expressed genes in in *Satb1^null^* EPCs were involved protein-protein interactions, cell signaling, and transcriptional regulation. These findings reconfirm previous reports on the role of SATB1 in different cell types and tissues^[3, 20, 21]^. Remarkably, IPA analysis revealed that SATB1-regulated genes in mouse EPCs are linked to functions that support the pluripotency of stem cells, facilitate self-renewal, and inhibit cell differentiation. This is in line with our previous findings in mouse and rat TS cells that SATB1 facilitates the proliferation of TS cells and inhibits trophoblast differentiation^[18, 27]^.

IPA analysis also revealed the differentially expressed genes in *Satb1^null^* EPCs linked to the WNT and TGFβ pathways, which are essential during the early stages of embryonic development. WNT and TGFβ signaling play crucial roles in trophoblast differentiation; overactivation of these pathways can lead to abnormal trophoblast cell development, which can affect the placenta^[28–30]^. An increased expression of key genes involved in the WNT signaling pathway like *Wnt2*, *Wnt3a*, *Wnt10a*, and *Wnt10b* in *Satb1^null^* EPCs might contribute to abnormal development of placenta. Similarly, BMP signaling has been reported to induce trophoblast development via N-cadherin via non-canonical ALK2/3/4-SMAD2/3-SMAD4 pathways^[31]^. It can also impact trophoblast differentiation both *in vivo* and *in vitro* via SMAD and TGF signaling^[32]^. TGFβ signaling acts via the SMAD pathway to induce differentiation of trophoblast cells^[33]^. Our data clearly showed exaggerated expression of *Bmp2*, *Bmp8*, *Smad1*, *Smad9*, and *Tgfbr2*. Thus, abnormal activation of the WNT, BMP, SMAD, and TGF pathways observed in *Satb1^null^* EPCs can ultimately result in defective placental development. UTF1 is a transcriptional repressor that interacts with the core pluripotency factors MYC and PRC2 to maintain self-renewal of stem cells^[34, 35]^. Thus, downregulation of *Utf1* expression in *Satb1^null^* EPCs may lead to decreased proliferation of trophoblast cells during placental development.

The overlap between the known genes that regulate placental development and the differentially expressed genes identified in *Satb1^null^* EPCs including *Adamts3, Mmp14, Calcrl, Syna*, and *Cdkn1c* underscore the importance of SATB1 in placental development. Among this critical group of genes, *Adamts3* was reported to regulate placental angiogenesis^[36]^. *Mmp14*, which is expressed in both human and mouse placentas^[37–39]^, regulates trophoblast functions including release of soluble endoglin from syncytiotrophoblasts, which is a critical feature in preeclampsia^[38]^. *Mmp14* can promote differentiation of syncytiotrophoblasts in humans and formation of the labyrinth zone in mouse placentas^[39]^. Moreover, *Calcrl* is required for the functions of CGRP^[40, 41]^. CGRP can increase cAMP levels and induce differentiation of trophoblast cells^[42]^. *Syna* is a retroviral envelope gene expressed in trophoblast cells and is involved in syncytiotrophoblast formation through fusogenic activity. *Syna* also plays a role in placental angiogenesis and is involved in the pathogenesis of preeclampsia^[43]^. *Cdkn1c* inhibits *Cdk1* and plays an important role in endoreduplication of genomic DNA during trophoblast differentiation to giant cells^[44]^. *Cdkn1c* maintains the integrity of the maternal-fetal surface and is involved in the allocation of maternal resources via the placenta^[45]^. Thus, SATB1 is found to regulate key genes that regulate trophoblast differentiation and the development of placenta.

## ACKNOWLEDGEMENTS

This study was supported by the funding from NIH/NICHD grant HD079363.

## CONFLICT OF INTEREST

The authors do not have any conflicts of interest.

## Notes

### Competing Interest Statement

The authors have declared no competing interest.

