## Supplementary figures and images for "SATB homeobox 1 regulated genes in the mouse ectoplacental cone are important for placental development"

### Supplemental Figure 1

#### Supplementary Figure 1

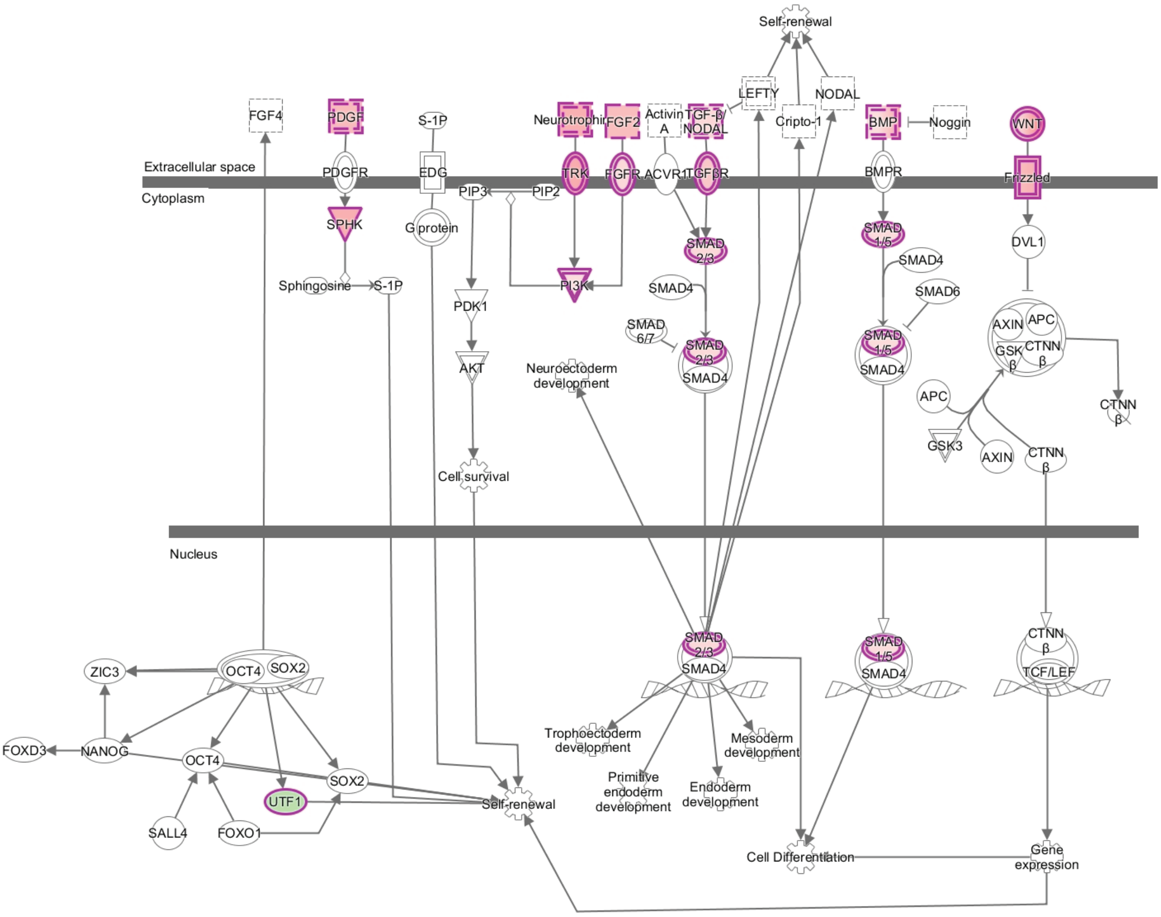
