## Supplemental Table 1 for "SATB homeobox 1 regulated genes in the mouse ectoplacental cone are important for placental development"

**Supplementary Table 1. List of primers used in the qPCR assays**

| Symbol | Reference mRNA | Forward Primer | Reverse Primer | Amplicon |
|---|---|---|---|---|
| *Wnt2* | NM_023653.5 | 192F-CTCCCTCTGCTCTTGACCTG | 293R-TGGCACATTGTCACACATCA | 102bp |
| *Wnt3a* | NM_009522.2 | 391F- CCATCTTTGGCCCTGTTCT | 477R-TCACTGCGAAAGCTACTCCA | 87bp |
| *Wnt10a* | NM_009518.2 | 221F- GGCGCTCCTGTTCTTCCTAC | 418R- ATGCCCTGGATAGCAGAGG | 198bp |
| *Wnt10b* | NM_011718.2 | 86F-TCACTCCCTCCCTTTTACCC | 404R-CTGAACAAAGCCAAGAACAGG | 319bp |
| *Bmp2* | NM_007553.3 | 1176F-GACTGCGGTCTCCTAAAGGTC | 1373R- CTGAGCAGCCTCAACTCAAA | 198bp |
| *Bmp8a* | NM_001256019.1 | 777F-ACCTATACCACGCCATGACC | 946R-AGCAGGGATCTGGGTTAGGT | 170bp |
| *Smad1* | NM_008539.4 | 961F-ACCTACCCTCACTCCCCAAC | 1153-CGTAAGCAACTGCCTGAACA | 193bp |
| *Smad9* | NM_019483.5 | 705F-ACCATTACCGCAGAGTGGAG | 786R-AGGCTGAGCTGAGGGTTGTA | 82bp |
| *Tgfbr2* | NM_009371.3 | 359F-GCTGCATATCGTCCTGTGG | 490R-TTCAGTGGATGGATGGTCCT | 132bp |
| *Utf1* | NM_009482.2 | 531F-GTCCCTCTCCGCGTTAGC | 705R-CAGAGTGTCGGTGCTCGTAA | 175bp |
| *Adamts3* | NM_001081401.2 | 911F-CAGGAACCTCTGTTGCCATT | 1083R-TACATCTCTGGGAGGCTGCT | 173bp |
| *Mmp14* | NM_177354.4 | 465F-CCGCCATGCAAAAGTTCTAT | 582R-GCCTTGATCTCAGTCCCAAA | 118bp |
| *Calcrl* | NM_018782.2 | 198F-GGCCCTATGATGATACAGCAA | 397R-AAGCAACCTGTGACCTTGGA | 200bp |
| *Syna* | NM_001013751.2 | 7198F-GCTGATTCATGGGAAAGGAA | 7247R-AGGCCCCTAACAGTGGTTCT | 50bp |
| *Cdkn1c* | NM_001161624.1 | 1013F-AGGAGCAGGACGAGAATCAA | 1202R-ACGTTTGGAGAGGGACACC | 190bp |
| *Rn18s* | NR_046237.1 | 1622F-GCAATTATTCCCCATGAACG | 1744R GGCCTCACTAAACCATCCAA | 123 bp |
